# Bacterial mechanosensing of surface stiffness promotes signaling and growth leading to biofilm formation by *Pseudomonas aeruginosa*

**DOI:** 10.1101/2023.01.26.525810

**Authors:** Liyun Wang, Yu-Chern Wong, Joshua M. Correira, Megan Wancura, Chris J Geiger, Shanice S Webster, Benjamin J. Butler, George A. O’Toole, Richard M. Langford, Katherine A. Brown, Berkin Dortdivanlioglu, Lauren Webb, Elizabeth Cosgriff-Hernandez, Vernita D. Gordon

## Abstract

The attachment of bacteria onto a surface, consequent signaling, and the accumulation and growth of the surface-bound bacterial population are key initial steps in the formation of pathogenic biofilms. While recent reports have hinted that the stiffness of a surface may affect the accumulation of bacteria on that surface, the processes that underlie bacterial perception of and response to surface stiffness are unknown. Furthermore, whether, and how, the surface stiffness impacts biofilm development, after initial accumulation, is not known. We use thin and thick hydrogels to create stiff and soft composite materials, respectively, with the same surface chemistry. Using quantitative microscopy, we find that the accumulation, motility, and growth of the opportunistic human pathogen *Pseudomonas aeruginosa* respond to surface stiffness, and that these are linked through cyclic-di-GMP signaling that depends on surface stiffness. The mechanical cue stemming from surface stiffness is elucidated using finite-element modeling combined with experiments - adhesion to stiffer surfaces results in greater changes in mechanical stress and strain in the bacterial envelope than does adhesion to softer surfaces with identical surface chemistry. The cell-surface-exposed protein PilY1 acts as a mechanosensor, that upon surface engagement, results in higher cyclic-di-GMP levels, lower motility, and greater accumulation on stiffer surfaces. PilY1 impacts the biofilm lag phase, which is extended for bacteria attaching to stiffer surfaces. This study shows clear evidence that bacteria actively respond to different stiffness of surfaces where they adhere *via* perceiving varied mechanical stress and strain upon surface engagement.

**Importance:** Bacteria colonize many types of biological and medical surfaces with a large range of stiffnesses. Colonization leads to the formation of biofilms, which cause costly and life-impairing chronic infections. However, whether and how bacteria can sense and respond to the mechanical cue provided by surface stiffness has remained unknown. We find that bacteria do indeed respond to surface stiffness in a way that is both consistent with expectations based on equilibrium continuum mechanics and that quantitatively impacts multiple aspects of early biofilm formation. This is a new understanding for the nascent field of bacterial mechanobiology. Furthermore, this finding suggests the possibility of a new category of approaches to hindering biofilm development by tuning the mechanical properties of biomedical surfaces.

## Introduction

Mechanosensing, including but not limited to responding to surface stiffness, is well-established to be an important cellular function in eukaryotes (1, 2). Much less is known about mechanosensing by prokaryotes (3–5). A few recent studies have shown that during biofilm formation, bacteria can sense and respond to mechanical cues, such as those arising from contacting a surface (6–12) and varying fluid flow over surface-bound bacteria (13, 14). For the biofilm-forming pathogen *Pseudomonas aeruginosa*, its cell-surface-exposed protein PilY1 has been proposed as a possible mechanosensor of surface adhesion (10, 13) and fluid shear (13). PilY1 is localized at the outer membrane (9, 10) and likely found at the tip of type-IV pili (TFP) as well (15). The extension and retraction of TFP power the twitching motility of *P. aeruginosa* on surfaces and are suggested to contribute to bacterial mechanosensing of surfaces (8, 11) and fluid shear (13).

*In vivo*, bacteria can experience a wide range of surface stiffnesses, from ultrasoft (dermal fillers have elastic moduli 0.02-3 kPa and living tissues 0.2-30 kPa) to hard (orthopedic implants have elastic moduli 5-300 GPa) (16, 17). In such diverse settings, biofilm formation commonly causes chronic infection, resulting in prolonged illness and high medical costs (18–20). Some research indicating that bacteria may be capable of sensing surface stiffness has recently emerged, showing that the initial accumulation of bacteria varied on surfaces of different stiffness (21–24). However, these earlier studies varied stiffness by varying characteristics such as cross-linking density or polymer concentration, which could also affect surface porosity or the density of attachment sites – in essence, changing at least two variables of the surface encountered. Furthermore, an inappropriate fabrication of surfaces with different stiffness may introduce unintended changes to other surface properties, such as adhesivity (*SI discussion*). Perhaps as a result, the literature on the effect of surface stiffness on bacterial accumulation on surfaces does not show consistent trends (21, 22, 24). Furthermore, there is currently very little understanding of how bacteria perceive the stiffness of surfaces to which they attach and how this input signal might allow the microbes to modulate their post-attachment accumulation accordingly. In addition, whether bacteria distinguish and respond to surface stiffness in later, post-accumulation stages of biofilm formation are also unknown. These knowledge gaps are of critical importance because they prevent the design of strategies for controlling biofilm development by manipulating surface stiffness.

To address these knowledge gaps, in the present study, we used thin and thick hydrogels coated on glass coverslips to create stiff and soft composite materials with the same surface chemistry, but different effective stiffnesses, and monitored *P. aeruginosa* through the early stages of biofilm formation on these surfaces. For surfaces exposed to a suspension of bacteria for one hour, we used quantitative microscopy to measure bacterial accumulation on materials of different effective stiffnesses. For the immediately-following stages of biofilm formation, characterized by bacteria reproducing on a surface rather than accumulating on the surface out of a suspended (planktonic) population, we measured the duration and growth rate of the biofilm lag phase and exponential growth phase, respectively.

For both the accumulation and reproduction stages of biofilm development, we show that bacteria actively recognize and respond to surface stiffness. When bacteria initially attach to a surface, both finite element modeling and experimental measurements of the activity of mechanosensitive ion channels show that attachment to stiff surfaces causes greater changes in the mechanical stress and strain state of the bacterial cell envelopes than does attachment to soft surfaces. PilY1 acts as a mechanosensor to transduce mechanical changes in the bacterial envelope into different intracellular levels of the second messenger cyclic-di-GMP (c-di-GMP). Using a proxy reporter for c-di-GMP levels, we measure higher levels of PilY1-dependent c-di-GMP production on stiffer surfaces than on softer. Higher levels of c-di-GMP lead to greater reduction in motility, a reduced likelihood of detachment, and, as a result, greater accumulation on the surface. Once the initial accumulation stage has passed, higher levels of cyclic-di-GMP are associated with a longer biofilm lag phase on stiffer surfaces.

## Results

### PilY1 allows *P. aeruginosa* to differentially accumulate on substrates of different stiffness

To eliminate effects arising from physicochemical properties of surfaces other than stiffness, such as adhesivity or porosity, we fabricated thin and thick hydrogels atop glass coverslips. (Fig. 1*A* and Fig. S1 *A*). Different hydrogel thickness had the same chemical compositions and the fabrication methods used did not alter the surface chemistry or topography (Fig. S1 *B* and *C*, Fig. S1 *F* and *G*). We chose this geometry-based approach to modifying substrate stiffness to avoid inadvertently altering material adhesivity along with stiffness, which has been observed before (*SI discussion*) - for instance, poly(dimethylsiloxane) (PDMS) can have different surface adhesivities associated with different stiffnesses, shown by polymer beads found to accumulate more on soft PDMS than on stiff PDMS (25).

**Fig. 1.**
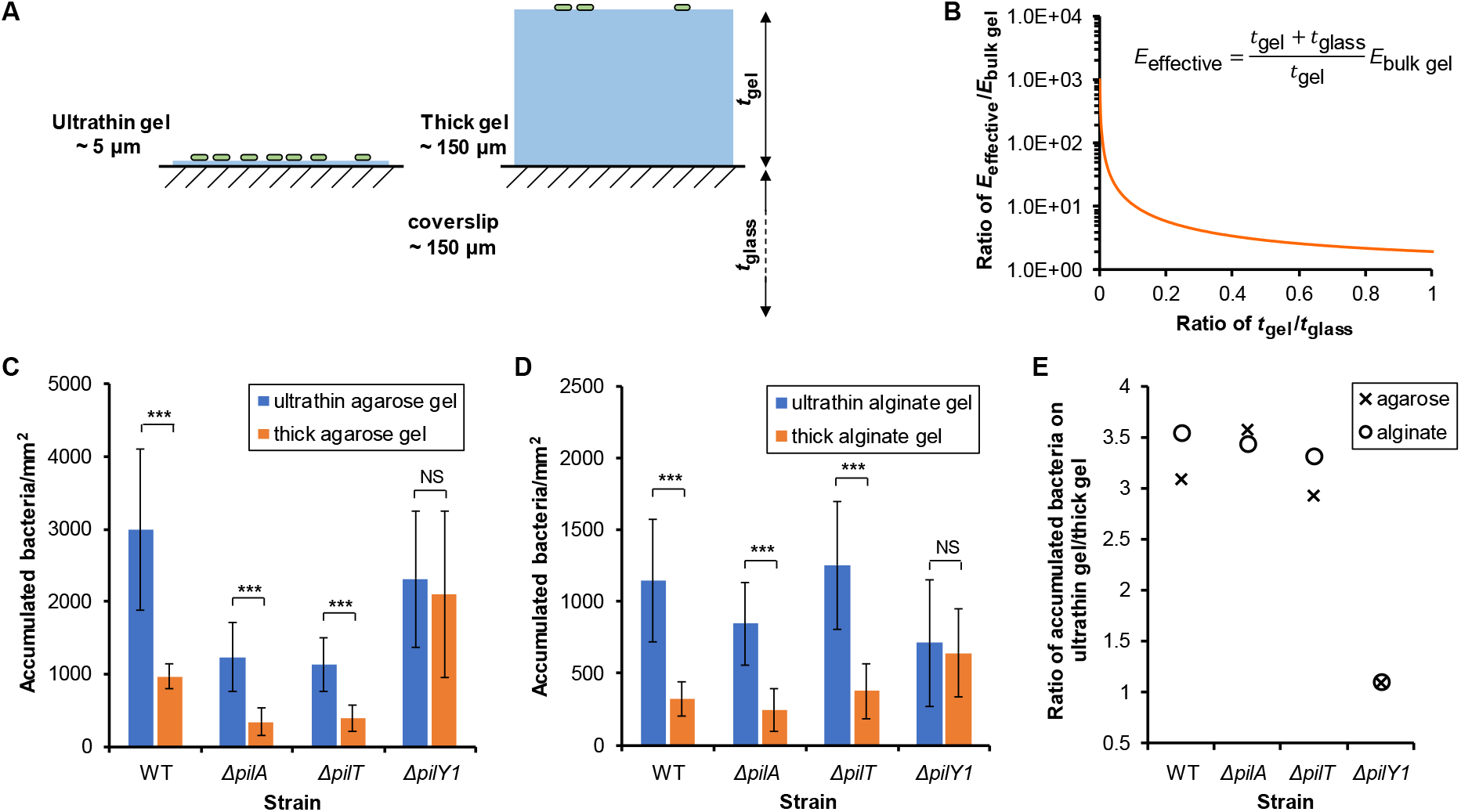
More bacteria accumulate on stiffer surfaces during one hour’s incubation for initial attachment. *(A)* Schematic illustration of composites with different thicknesses of hydrogel, *t*_gel_, on top of glass coverslips with constant thickness *t*_glass_. *(B)* The effective Young’s modulus of the hydrogel-coverslip composite (*E*_effective_), where *E*_bulk gel_ is the modulus of bulk hydrogel. *(C* and *D)* The accumulation of WT, Δ*pilA*, Δ*pilT* and Δ*pilY1* on thin and thick hydrogel composites after incubating with surfaces for one hour. Data are means ± SD. \*\*\**P* <0.001; NS, not significant (*P* = 0.28 for agarose; *P* = 0.29 for alginate); analysis of variance (ANOVA) test. *(E)* The ratio of accumulated bacteria on thin to that on thick hydrogel composites.

To check that the composite materials we made had the same surface adhesivity regardless of gel thickness, we incubated both thin and thick hydrogel composites with a suspension of fluorescent polystyrene polymer beads for one hour, and imaged the number of beads attached using confocal microscopy. We verified that the numbers of polystyrene beads that attached did not significantly differ with hydrogel thickness (*SI Discussion*, Fig. S1 *D* and *E*). Thus, we conclude that hydrogel thickness does not impact passive physicochemical adhesion to surfaces.

However, the thickness of the hydrogel coated onto rigid glass coverslips does impact the stiffness of the resulting composite material. Linear elasticity theory was used to derive a closed-form expression for the effective elastic modulus (*E*_effective_) of hydrogel-coverslip composites (*SI Discussion* and Equation. S7). For a hydrogel thickness (*t*_gel_) comparable to the 1 micron size of a bacterium, the *E*_effective_ increases sharply with decreasing *t*_gel_ (Fig. 1*B*). According to this model, the composites with thin (~5 μm) hydrogels are approximately 16 times stiffer than those with thick (~150 μm) (Fig. 1*B* and Table S1). We also used a nanoindenter to experimentally impose loads on both types of composites that achieved indentation depth comparable to what bacteria realistically experience in the experiments (Fig. S1 *H*). The indentation for a given force and tip geometry was consistently less for the thin gel than for the thick gel, thereby validating that the composite made with thin gel is stiffer than the composite made with thick (*SI Discussion*, Fig. S1 *H*).

To assess the impact of surface stiffness on the accumulation of bacteria on the surface, we incubated the bacterial suspension for one hour with hydrogel-coverslip composites and measured the bacterial accumulation on these surfaces by visualizing the number of bacteria using phase contrast microscopy. Consistent with some previous reports (21–23) but not with others (24), wild type *P. aeruginosa* cells (WT) accumulated significantly more on stiffer composites than on softer (Fig. 1 *C* and *D*), as did mutants without TFP (Δ*pilA*) and mutants without the PilT retraction motor (Δ*pilT*) (Fig. 1 *C* and *D*), by a factor of ~3.3 for all three strains (Fig. 1*E*). Thus, while functional TFP can increase the “baseline” accumulation, they have no measurable impact on the greater likelihood of accumulating on stiffer surfaces. Similar effects were found for the more starkly-contrasting case of glass versus agarose gel surfaces, (*SI Discussion*, Fig. S2 *A*).

In contrast, mutants without PilY1 (Δ*pilY1*) accumulated equally on effectively-stiff and effectively-soft composites (Fig. 1 *C and D*). This indicates that *P. aeruginosa* requires the cell-surface-exposed protein PilY1 for distinguishing between, and responding to, different surface stiffnesses. Again, similar effects were found for the more starkly-contrasting case of glass versus agarose gel surfaces, implying a much more muted response to stiffness difference by Δ*pilY1* (*SI Discussion*, Fig. S2 *A*).

Adhesive forces will tend to increase the area of the bacterium in contact with the surface, by deforming the bacterium and the surface. The energy costs for deforming the bacterium and the surface will depend on the elasticity of each. Mechanical equilibrium will be found by minimizing the sum of elastic energy costs (from cell and substrate deformation) and the adhesive energy benefit (from contacting area). Therefore, for constant adhesive area and bacterial elasticity, we expect that the deformation of the bacterial cell envelope will depend on the elasticity of the substrate. In *Escherichia coli*, cell membrane proteins can sense and respond to changes in mechanical stresses in the bacterial cell envelope resulting from surface attachment (3, 26, 27). We hypothesized that, upon adhesion to surfaces of different stiffnesses, *P. aeruginosa* should undergo different changes in mechanical stress and strain upon surface engagement, which might be perceived by the cell-surface-exposed protein PilY1. Notably, this type of mechanical stress is distinct from biological stress, involving unfolded or misregulated proteins in the cell envelope, which has also been shown to relate to surface sensing in *P. aeruginosa* (28).

### Stiffer substrates lead to greater changes in mechanical stress and strain in the bacterial cell envelope

To elucidate the relationship between surface stiffness and mechanical stresses in adhering bacteria, we developed finite element models (Fig. 2*A*, and Fig. S3) to simulate bacterial attachment to gel-coverslip composites. At the molecular level, bacterial surface properties and how they impact attachment to substrates are complex and not well-known (29). Therefore, we approximated the adhesion process by displacing bacteria into contact with surfaces (*SI Discussion*, Fig. S3 *C* and *D*). Using our models, we compared the bacterial envelope mechanics for bacteria interacting with stiff and soft surfaces for a range of contact-increasing displacements. For any given displacement, the total contact area is greater for bacteria on a soft surface than on a stiff one (Fig. 2*B*), reflecting the fact that the energy cost for deforming a soft material is lower than the cost for deforming a stiff one by the same amount. The initial, free-floating cells were subjected only to a turgor pressure (biologically, this arises from the osmolarity difference between the cytoplasm and the exterior), so that bacteria were in a pre-stressed state. Contact with a surface leads to a decrease in membrane stresses on the outer membrane, an increase in circumferential strain on the inner membrane, and the development of contact pressure (Fig. 2*C*, Fig. S4 *A* to *I*). These changes are all more pronounced when the surface is stiffer.

**Fig. 2.**
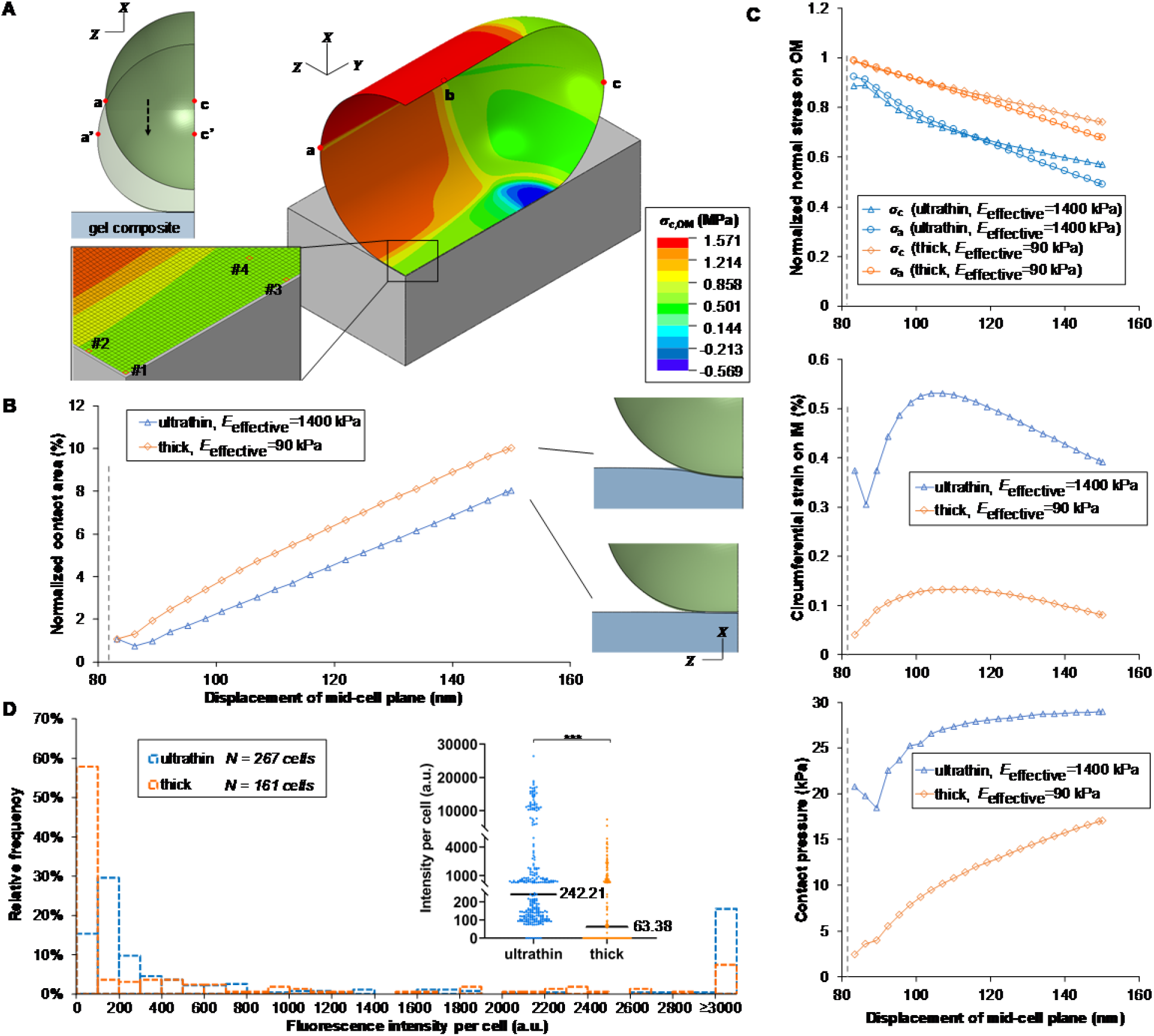
Adhesion to a stiffer surface leads to greater changes in mechanical stress/strain in the bacterial envelope and increased permeation of the bacterial cell membrane by sodium. *(A)* The finite element model and schematic illustration. Displacement along −X coordinate is applied on curve abc to bring the cell into contact with the surface. The heat map denotes the circumferential stress on OM (outer membrane). Note that the stress result on the substrate is not shown here. Inset: The representative elements analyzed in this study. *(B)* Contact area with different degree of indentation (displacement along −X coordinate). Contact area is normalized to the cellular surface area in the undeformed configuration. The dash line denotes when the cell first contacts the substrate. *(C)* OM stresses become less tensile whereas IM (inner membrane) strain increases at element #1 upon surface adhesion. The degree of changes is greater on a stiffer substrate. Contact pressure is greater on a stiffer substrate. Subscript c denotes the circumferential direction and subscript a denotes the axial direction. Stresses are normalized to their respective values during the free-floating state and strains are the net change with respect to their respective values during the free-floating state. *(D)* The histogram shows the average intracellular fluorescence intensity per cell of attached WT on thin and thick agarose gel composites after incubating with surfaces for one hour. Inset: Dot plot of the histogram, shown with median values. \*\*\**P* <0.001; Mann-Whitney u test. This indicates a statistically-significant difference between fluorescence intensity distributions and between median fluorescent intensities for cells on thin and thick gel composites.

To validate the trends shown by our modeling results, we compared the membrane tension in bacteria attached to gel-coverslip composites of different effective stiffnesses, by comparing the activity of mechanosensitive ion channels. These channels are located on the inner, cytoplasmic membrane (30) and act as transducers of membrane tension - closed when the membrane is at low tension and open when the membrane is at high tension, allowing ions to pass through (30, 31). The two major mechanosensitive ion channels are large-conductance- and small-conductance- (MscL- and MscS-, respectively) type channels. When open under increased membrane tension, these channels provide non-selective pores of large and small diameter, respectively, through which sodium ions, Na^+^, can pass in very similar ways (31). We pre-loaded bacteria with a fluorescent indicator for Na^+^ and then allowed them to sit for one hour attached to thin and thick agarose gels, in the presence of excess external Na^+^, before measuring the indicator brightness distribution as a proxy for internal Na^+^ levels.

The brightness distribution for bacteria on stiff substrates had a peak at 100-200 arbitrary units (a.u.), whereas the brightness distribution for bacteria on soft substrates had a peak, representing more than 60% of cells, at 0 to 100 a.u. (Fig. 2*D*). Both the median (Fig. 2*D* inset) and the mean fluorescence intensity of bacteria on stiff substrates were significantly greater than that of cells on soft substrates – cells on stiff substrates had a mean fluorescence intensity of 2840.70 a.u. [2217.75 a.u., 3463.65 a.u.] (95% confidence interval) and cells on soft substrates had a mean fluorescence intensity of 677.97 a.u. [478.28 a.u., 877.67 a.u.] (95% confidence interval). These results show that bacteria on stiff substrates are more permeable to Na^+^ than are bacteria on soft substrates. Since mechanosensitive ion channels increase permeability upon increased membrane tension, we interpret this finding as indicating that bacteria have higher membrane tension when attached to stiffer materials.

Thus, these experimental results are consistent with our modeling results and therefore support the idea that the assumptions underlying our modeling are reasonable in this regime.

<u>Adhesion-induced changes can only happen following, not preceding, bacterial contact with surfaces.</u> Since gel thickness does not impact physiochemical surface adhesivity (Fig. S1 *D* and *E*), we expect bacteria to have equal likelihood of encountering and initially sticking to stiff and soft substrates. Therefore, this is the first report, to our best knowledge, showing that greater accumulation on stiffer composites must arise as the result of something that happens after initial surface engagement – i.e., *an active bacterial response to substrate stiffness*. We hypothesized that WT initially adhered to thick hydrogels will be more likely to detach than cells initially adhered to thin hydrogels and that this difference should require PilY1.

### PilY1 transduces substrate stiffness to adjust flagellar spinning and the detachment rate

Shortly after encountering a surface, many *P. aeruginosa* cells are reversibly tethered by their flagella, which drive in-place spinning (32); spinning facilitates detachment from surfaces (33, 34). Deficiency in spinning is associated with decreased probability of detachment (33). Bacteria can also use TFP to move laterally on surfaces, but, during the first hour after bacteria were introduced to hydrogels (i.e., what we have termed the accumulation stage), we found that the vast majority of surface motility was in the form of spinning (Fig. S5 *A* and *B*). Therefore, we tracked the center-of-mass speed of surface-adhered bacteria (Fig. 3 *A-D*) as for a measure of spinning motility. We expect that a population with faster-spinning bacteria will have a higher rate of detachment (35).

**Fig. 3.**
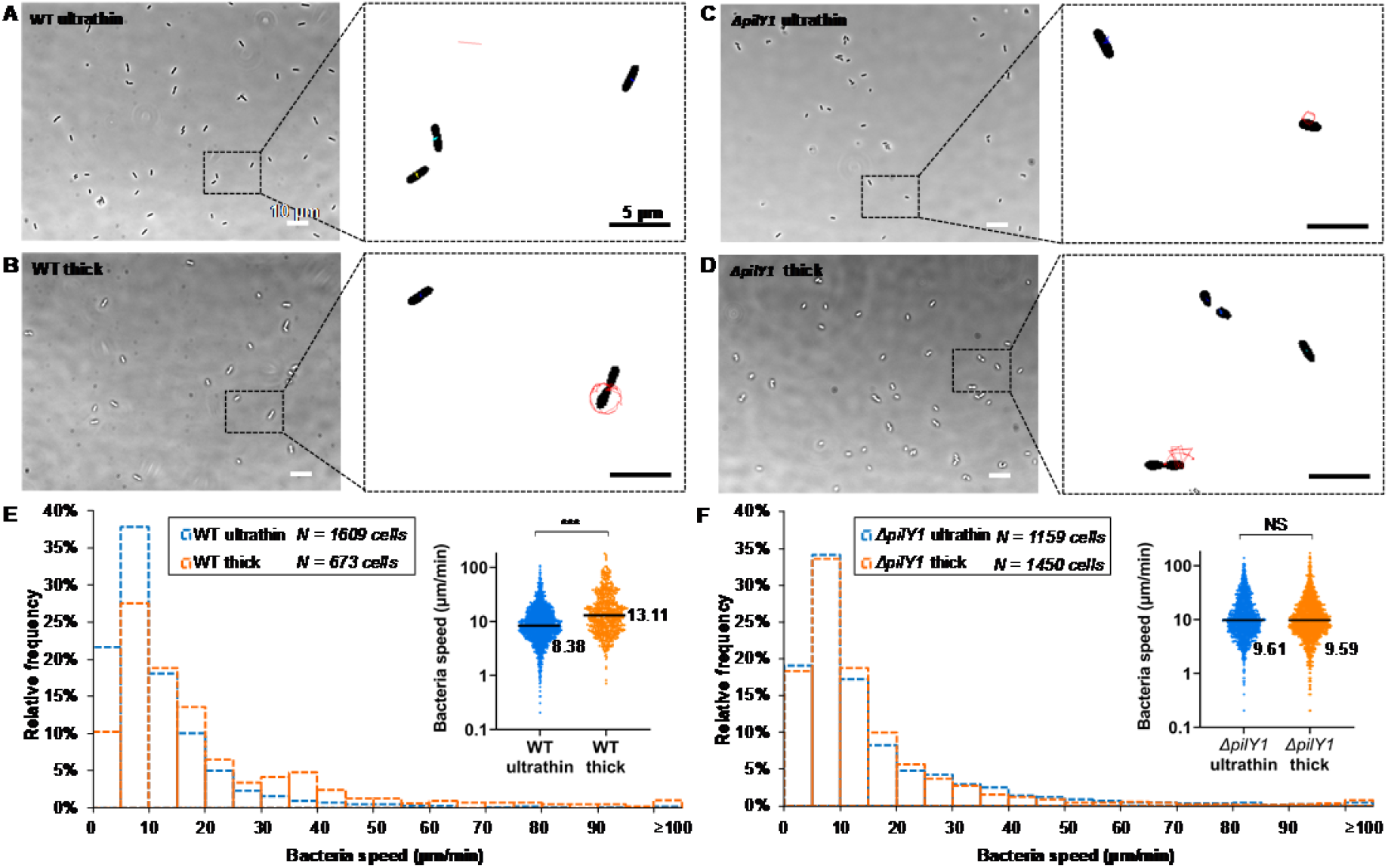
Adhered bacteria spin during the first hour of accumulation. *(A-D)*, Phase contrast images of WT and the Δ*pilY1* mutant adhered to thin and thick agarose gel composites. Insets: Tracked trajectories of bacterial centers-of-mass over 62.6 s. *(E* and *F)*, Histograms showing speed distributions of WT and the Δ*pilY1* mutant on thin and thick gel composites. Insets: Dot plots of the corresponding histogram. The median value is written to the right of each plot. \*\*\**P* <0.001; Mann-Whitney u test. *** indicates a statistically-significant difference in the distributions of WT speeds on thin and on thick gel composites and that the median speed of WT adhered to thick gel composites was higher, with statistical significance, than that of WT to thin gel composites. In contrast, NS (not significant) indicates that there is no statistically-significant difference in the distributions of speeds or in the median speeds of the Δ*pilY1* mutant on the two composite types (*P* = 0.66, Mann-Whitney u test).

For WT, the distribution of spinning speeds on soft substrates was much broader than stiff substrates (Fig. 3*E*). Both the median (Fig. 3*E* inset) and the mean speeds on soft substrates were significantly higher than on stiff substrates - mean speed of 20.06 μm/min [18.43 μm/min, 21.68 μm/min] (95% confidence interval) on soft composites and mean speed of 11.46 μm/min [10.95 μm/min, 11.97 μm/min] (95% confidence interval) on stiff composites. In summary, WT are more likely to spin rapidly on soft substrates than on stiff substrates. Upon tracking cells, we indeed found that WT were significantly more likely to detach from soft substrates (30 detachment events among 673 tracked cells) than from stiff substrates (10 detachment events among 1609 tracked cells) (*P* <0.001, χ^2^ test) (Fig. S5 *C*). This is *an active bacterial response to substrate stiffness*.

For the Δ*pilY1* mutant, the peak spinning speed was unchanged from that of WT (Fig. 3 *E* and *F*), suggesting that loss of PilY1 does not intrinsically disrupt spinning motility. However, for the Δ*pilY1* mutant, neither the distributions of spinning speeds nor the median spinning speeds were significantly different on stiff and soft substrates (Fig. 3*F*). The mean speed was 15.08 μm/min [14.16 μm/min, 16.01 μm/min] (95% confidence interval) on thin gels and 14.86 μm/min [13.92 μm/min, 15.81 μm/min] (95% confidence interval) on thick. Furthermore, the Δ*pilY1* mutant was equally likely to detach from thin and thick gels (*P* = 0.78, χ^2^ test) (Fig. S5 *C*). These results are strikingly unlike those for WT, and imply that *P. aeruginosa* lacking PilY1 do not adjust their spinning motility, and therefore their likelihood of detachment, in response to surface stiffness.

From these findings, we infer that PilY1 is required for early sensing of substrate stiffness in *P. aeruginosa*, and PilY1 is linked to regulating flagellar activity either up or down - increasing spinning speed on soft surfaces and decreasing spinning speed on stiff surfaces (Fig. S5 *D* and *E*). Notably, we find a linear correlation between spinning speed and the probability of detachment (Fig. S5 *D* and *E*) This finding raises the question of what provides the *causative linkage* between PilY1 and changes in flagellar activity.

### Substrate stiffness impacts c-di-GMP signaling in a PilY1-dependent manner during accumulation

The intracellular second messenger c-di-GMP is broadly used by many bacteria to regulate many cellular processes, including the sessile-to-motile transition, biofilm formation, and flagella-mediated motility (36). Therefore, to see whether PilY1 modulates c-di-GMP dynamics in response to surface stiffness, we used a validated reporter plasmid, P_*cdrA*_∷*gfp*, that produces green fluorescent protein (GFP) in response to increases in c-di-GMP (37); this plasmid was previously used to study c-di-GMP signaling in bacterial mechanosensing of shear (13).

For bacteria containing PilY1, we found a sharp rise in c-di-GMP levels during the initial hour of accumulation (−1 to 0 h in Fig. 4 *A* and *C*), which is consistent with previous findings that, c-di-GMP levels in *P. aeruginosa* increase upon surface attachment (9, 13, 38). At the end of the “accumulation” hour (i.e., the beginning of the incubation time), WT on stiff substrates had significantly higher c-di-GMP levels than did WT on soft substrates. This meshes with our finding that WT on stiff substrates had lower spinning motility than those on soft substrates (Fig. 3*E*), as high levels of c-di-GMP inhibit bacterial motility (36). It also suggests that the causative linkage between PilY1 and changes in flagellar activity (which, in turn, modulate the likelihood of detaching from the surface), is through PilY1-controlled c-di-GMP signaling.

**Fig. 4.**
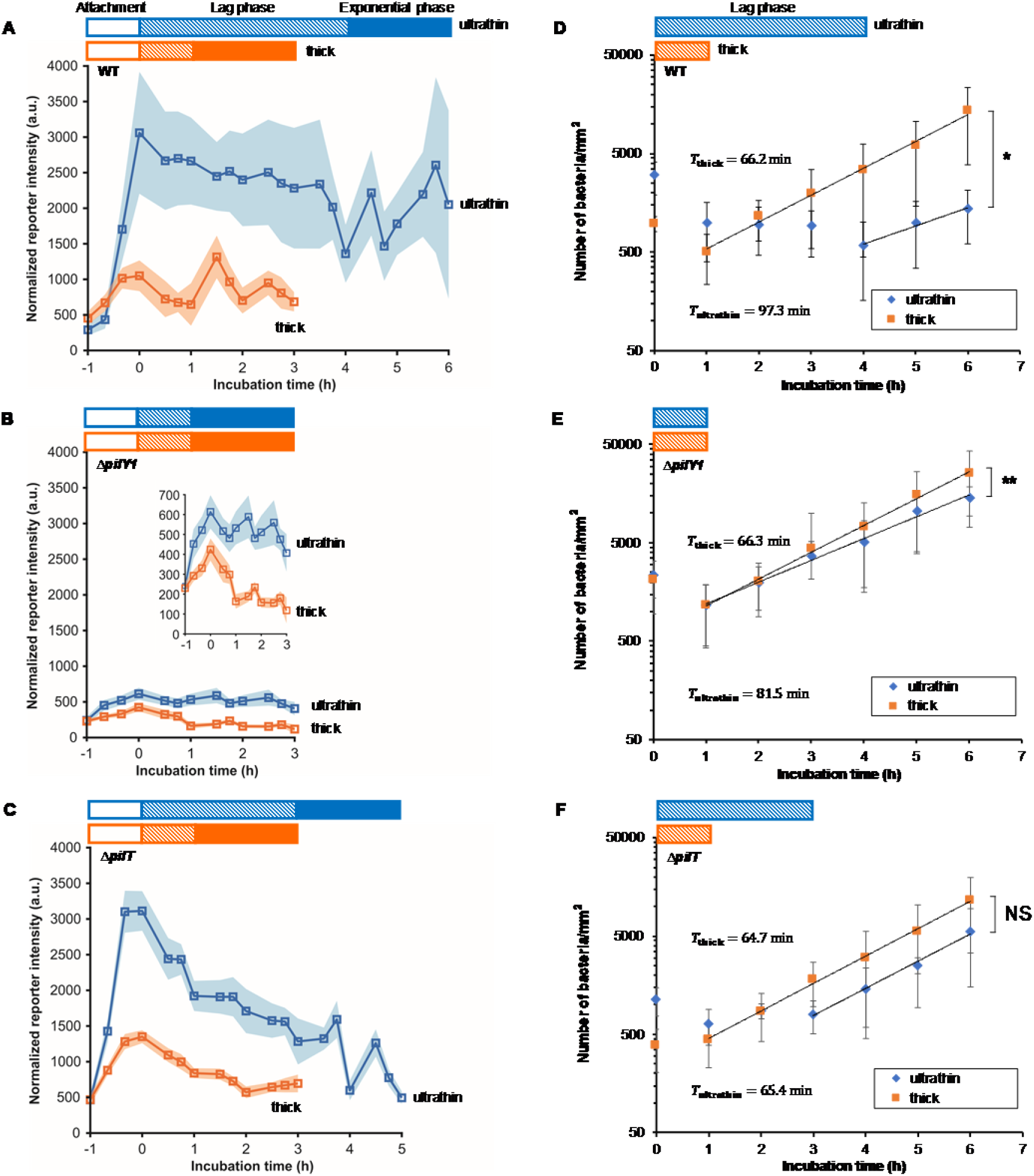
On surfaces with different stiffnesses, PilY1 acts to mediate the duration of the lag phase in biofilm growth and the levels of the intracellular signal c-di-GMP, and PilT is required to mediate the growth rate of the exponential phase of biofilm growth. *(A-C)*, Fluorescent reporter for changes in intracellular c-di-GMP in WT and the Δ*pilY1* and Δ*pilT* mutants during accumulation, lag phase, and exponential phase. The initial hour of accumulation on a surface is designated by −1 to 0 h, shown by hollow color bars. For each sample, exponential phase was observed for two hours, shown by solid color bars. Squares represent mean levels of c-di-GMP at each time point, linked by lines as a guide to the eye. Shaded regions correspond to 95% confidence intervals. The inset in (*E*) shows c-di-GMP reporter intensity in the Δ*pilY1* mutant with a smaller y-axis range. *(D-F)*, Growth dynamics of attached WT, and the Δ*pilY1* and Δ*pilT* mutants on thin and thick agarose gel composites. Data are means ± SD. The data at 0 time point corresponds to the end of one hour of bacterial accumulation on gel surfaces. Hatched color bars show the length of the lag phase. The doubling time, *T*, is calculated by the equation *T* = ln2/*α*, where *α* is the growth rate of bacteria on surfaces (equations of exponential regression, f(*t*) = A*e*^at^, where *t* is the incubation time). \*\**P* <0.01; \**P* <0.05; NS, not significant; analysis of covariance (ANCOVA) test. ** and * indicate that the growth rate *α*_thin_ is significantly different from *α*_thick_ for WT and for the Δ*pilY1* mutant, while NS means the difference in growth rates on thin and thick gel composites are not significant for Δ*pilT* (*P* > 0.1).

On both soft and stiff substrates, the Δ*pilY1* mutant had much lower c-di-GMP levels than did WT (Fig. 4 *A* and *B*); this is consistent with the role of PilY1 in regulating c-di-GMP production (9). The mean level of c-di-GMP at the end of the “accumulation” hour was ~2.9 times higher on stiff substrates than on soft for WT, while only ~1.4 times higher for the Δ*pilY1* mutant (Fig.4 *A* and *B*), consistent with a loss of the ability to discriminate surface stiffnesses. This finding is also consistent with the causative connection that PilY1 mechanosensing that controls c-di-GMP signaling levels is required for bacterial mechanoresponse to substrate stiffness during the initial “accumulation” phase.

This finding also raises the question of how PilY1 mechanosensing, and consequent changes in c-di-GMP signaling, impact the growth of the bacterial population on the surface.

### PilY1 impacts biofilm growth in the lag phase, in response to surface stiffness, by modulating c-di-GMP signaling

When planktonic bacteria are introduced into new liquid medium, they experience a temporary period of non-replication, termed the “lag phase” (39). After attachment to a glass surface, *P. aeruginosa* also undergoes a lag phase before exhibiting exponential growth (40, 41). However, unlike the planktonic lag phase, the lag phase of biofilm growth involves a combination of bacteria replication and detachment from surfaces, such that the population of surface-bound bacteria does not increase (40, 41).

After allowing bacterial accumulation on surfaces for one hour, we replaced the bacterial suspension with fresh, sterile culture medium. We designate this timepoint the beginning of the incubation time (0 h in Fig. 4). The duration of the lag phase, from the beginning of the incubation time to the onset of exponential growth, is given by the lag time, *τ*_lag_, indicated by hatched color bars in Fig. 4. WT populations had a *τ*_lag_ of 4 h on stiff substrates and 1 h on soft substrates, but Δ*pilY1* populations had the same *τ*_lag_ of 1 h on both substrate types (Fig. 4 *D* and *E*). Similar results were found for bacterial growth on bulk gels (soft) and glass slides (stiff) (Fig. S2 *B* and *C*). These results show that surface stiffness can markedly impact the growth of the sessile bacterial population, and that PilY1 is key for this process as well as for the accumulation preceding incubation. When PilY1 was complemented back on an arabinose-inducible plasmid, the Δ*pilY1* mutant populations again had different *τ*_lag_ on stiff and soft substrates (*SI Discussion*, Fig. S6 *C*), confirming PilY1’s role in surface stiffness sensing.

On both stiff and soft substrates, c-di-GMP levels in WT fell during the lag phase and subsequently oscillated once populations entered the exponential growth phase (Fig. 4*A*). The high level of c-di-GMP induced by the initial mechanical stimulus of surface contact (0 h in Fig. 4*A*) allows bacteria to sense the surface and initiate a sessile lifestyle. However, it would be a metabolic burden for cells to maintain such high c-di-GMP levels in the following biofilm development. We speculate that bacteria may have to decrease the c-di-GMP level to allow the beginning of exponential biofilm growth on surfaces; this speculation is consistent with the work of others (42, 43). If so, the longer *τ*_lag_ for WT on stiff surfaces than on soft likely arises from the much higher initial c-di-GMP levels on stiff surfaces and the consequent need for more time to gradually decrease c-di-GMP levels. For Δ*pilY1*, low initial levels of c-di-GMP are associated with a short *τ*_lag_ on both stiff and soft substrates (Fig.4 *B* and *E*).

We conclude that PilY1 is a required element for controlling *P. aeruginosa*’s initial c-di-GMP response to surface stiffness and consequent lag time in early biofilm growth.

## Discussion

The mechanical equilibrium of a system consisting of a bacterium adhering to a surface will be found when the net mechanical energy is minimized. Adhesion energy, which is energetically favorable and negative in sign, will increase in magnitude as the adhering area increases. The elastic energy costs for deforming the bacterium and the surface, to increase the adhering area, are energetically unfavorable and positive in sign. Since the elastic energy cost for a given deformation of a soft surface is less than the elastic energy cost for the same deformation of a stiff surface, we expect more of the elastic energy cost to be borne by the bacterium when the surface is stiff than when it is soft. Therefore, for surfaces that have the same adhesive properties, as for the thick and thin gel composites we use here, equilibrium mechanics leads us to expect bacteria adhering to soft surfaces will deform less than will bacteria adhering to stiffer surfaces; this has been confirmed by finite element modeling (Fig. S3 *A* and *B*). Thus, adhesion will result in greater changes in envelope stress for bacteria attached to stiff surfaces than for bacteria attached to soft surfaces. This expectation is supported by both finite element modeling and experiments measuring differences in the membrane tension as reflected by the opening of mechanosensitive ion channels in the inner membrane.

Our experimental results show that PilY1 is a key sensor that transduces mechanical changes upon surface engagement into c-di-GMP signaling. PilY1 is a surface-exposed protein found associated with the TFP tip (9), so PilY1 may be responding to the compressive loading incurred due to surface adhesion, a stress state identified in the modeling. A recent study suggested that the conformational changes of PilY1 lead to stimulation of bacterial c-di-GMP production and biofilm formation (44). The compressive loading may hence engender the required conformational changes on PilY1 for biofilm initiation. Modeling shows that bacteria adhered to stiffer surfaces will have a greater decrease in the tension in their outer membrane than will bacteria adhered to softer surfaces. Our experimental results show that the bacteria adhered to stiffer surfaces have higher c-di-GMP levels, resulting in lower spinning motility, less detachment, and greater accumulation in the first hour after exposure to the surface. Later, bacteria attached to stiffer surfaces need a longer lag phase in which to reduce their c-di-GMP level and begin exponential growth.

At this exponential-growth phase of biofilm development, our data suggests that the pilus retraction motor PilT may also be involved in responding to surface stiffness in a way that modulates c-di-GMP level and growth rate (Fig. 4 C and F); see *SI discussion*. The differential response of mechanosensitive ion channels to surface stiffness (Fig. 2 D) opens the possibility that mechanosensitive ion channels may play a role in the initial development in biofilms on surfaces, although we have not investigated that specifically.

From equilibrium mechanics, changes in bacterial stiffness would be expected to change the deformation in bacterial cells upon surface attachment, as stiffer bacteria would deform less and softer bacteria would deform more. This, in turn, would alter the mechanosensing response to surface attachment. It has recently been shown that *P. aeruginosa* maintain tight genomic control of their stiffness (45). This clearly has benefits for protecting the bacteria against mechanical stress, such as osmotic pressure; our work suggests that this may also benefit bacteria by safeguarding the surface-sensing response, which is essential to this biofilm-former.

Finally, our study has implications for what types of bacteria are likely to mechanosense surface contact through an envelope protein. The effective modulus of our composites with thin gel was roughly 1 MPa and the effective modulus of composites with thick gel was slightly less than 100 kPa (Table S1). These values bracket the stiffnesses reported for *P. aeruginosa* and other Gram-negative bacteria (46–48). Bacteria themselves are a composite material, comprising the softer cytoplasmic interior and the stiffer envelope. The Young’s modulus for the envelope material *per se* of Gram-negative bacteria is roughly several tens of MPa, and the envelope material of Gram-positive bacteria probably has a similar modulus (49, 50). Our finite element modeling identifies bending as the major envelope deformation modality in the contact zone as bacteria attach to surface. According to the Kirchhoff-Love plate theory (51), the flexural rigidity of a thin plate (effectively the modulus that measures the energy cost for bending a plate) is characterized by *Et*^3^ ⁄ 12(1 − *ν*^2^) ∝ *t*^3^, where *E* is the Young’s modulus of the plate, *ν* is the Poisson’s ratio, and *t* is the plate thickness. Gram-negative bacteria have a much thinner peptidoglycan cell wall than do Gram-positive bacteria (the cell wall of *P. aeruginosa* (Gram-negative) is ~3 nm thick (52) and that of *B. subtilis* (Gram-positive) is ~30 nm thick (53)). Therefore, it is unsurprising that whole cells of Gram-positive bacteria appear to be stiffer (54) than Gram-negative bacteria (49, 55–57). This suggests that Gram-positive bacteria will deform less than will Gram-negative bacteria upon adhesion to the same surface, because the energetic cost for deforming Gram-positive bacteria will be higher, and therefore that Gram-positive bacteria may be less well-adapted to using surface proteins to sense and respond to surface stiffness. This inference is in agreement with previous reports that Gram-positive bacteria do not respond to surface stiffness in the same way as Gram-negative bacteria (58, 59).

In conclusion, the work presented here provides a new understanding of bacterial response to surface mechanics in early biofilm development, which may point the way to new approaches to control biofilm infection on devices by manipulating the mechanical properties of materials. In the current state of the art, antimicrobial coatings have been developed that kill bacteria (60, 61) and antifouling surfaces have been developed that resist the attachment of bacteria (60–64). Manipulating mechanical properties to hinder biofilm development has not been explored, and materials with biofilm-hindering mechanics could also incorporate antimicrobial or antifouling properties, potentially to synergistic effects.

Our study may also suggest how different environmental mechanics *in vivo* could affect the course of infection. Examples of important sites of biofilm infection where mechanics can vary include airway mucus in Cystic Fibrosis patients, which is stiffer and more amenable to biofilm formation than that of healthy people (65). Wounds that are kept moist are less susceptible to infection and have better healing than wounds that are allowed to dry (66); drying, by reducing water content, will act to increase stiffness.

## Materials and Methods

We used *P. aeruginosa* PAO1 WT and Δ*pilA*, Δ*pilT*, Δ*pilY1* mutants (67). Studies of bacterial accumulation, motility, growth, and c-di-GMP production were done with bacteria that contained the plasmid P_CdrA_∷*gfp*. This plasmid is a verified reporter for cyclic-di-GMP (c-di-GMP); the *cdrA* gene, and thus green fluorescent protein (GFP), is upregulated by c-di-GMP level (37). Strains containing a promotorless control plasmid pMH487 were used to measure background GFP expression independent of c-di-GMP levels (13). Strains with the P_CdrA_∷*gfp* and pMH487 plasmids were grown with 60 μg/mL Gentamycin (Sigma-Aldrich, G1914) for plasmid selection. Fluorescence measurements using the sodium ion indicator were performed using WT that did not contain any plasmid. Details of fabrication of hydrogel composites, microscopy measurement of bacterial accumulation, biofilm growth on surfaces and c-di-GMP signaling, finite element modeling of the cell-surface interaction, intracellular level of sodium ions in surface-adhered bacteria, tracking motility of surface-adhered bacteria, and image and data analysis are described in *SI Materials and Methods*.

## Acknowledgments

This work was supported by grants from the Cystic Fibrosis Foundation (Gordon 201602808-001), the National Science Foundation (NSF) (727544 and 2150878, BMMB, CMMI), and the National Institutes of Health (NIH) (1R01AI121500-01A1, NIAID), all to Vernita Gordon. Additional support was provided through the NSF (2119716, DMREF, CMMI) to Berkin Dortdivanlioglu, through the NSF (1807215 and 22032414, CHE, MPS) to Lauren Webb, and the NIH (R37 AI83256) to George O’Toole.

## Legends of figures and tables in the supplemental material

**Fig. S1.** *(A)* Schematic of setup used to measure gel thickness by microscopy. Changing hydrogel thickness does not alter surface chemistry or passive physicochemical adhesion to the surface. FTIR spectra of *(B)* agarose and *(C)* alginate gel composites with two thicknesses. One spectrum from each sample is shown here. The dash-dot lines indicate the location of characteristic peaks. The number of beads attached on agarose gel composites *(D)* and alginate gel composites *(E)* after incubation with bead suspension in NaCl buffer for 1 h. NS, not significant (*P* = 0.15 for agarose; *P* = 0.62 for alginate); ANOVA test. NS indicates that the attachment of beads on thin and on thick gels are not significantly different for agarose gel composites and for alginate gel composites. A collection of representative images showing the surface of *(F)* ‘thin’ and *(G)* ‘thick’ hydrogels as visualized by Cryo Surface Electron Microscopy. It seems not possible to detect a difference between the surface structure of the thin and thick gels. *(H)* Nanoindentation results of 3% (w/w) agarose gel samples. The relation between maximum load and maximum indentation of indentation curves of gels subjected to large indentation. Data are means ± SD.

**Fig. S2.** *(A)* One hour after introducing bacterial suspension to surfaces, the numbers of WT and the Δ*fliC*, Δ*pilA*, and Δ*pilY1* mutants on glass surfaces are respectively 16.8, 20.5, 15.8 and 3.5 times higher than those on bulk agarose gel surfaces. NS, not significant (*P* = 0.36 for WT vs. Δ*fliC*; *P* = 0.63 for WT vs. Δ*pilA*; χ^2^ test). NS indicates that the ratio of accumulated mutant bacteria on glass and agarose is not significantly different from that for WT. \*\*\**P* <0.001; χ^2^ test. *** indicates the ratio of accumulated the Δ*pilY1* mutant on glass and agarose is significantly smaller than that for WT. Growth curves of WT *(B)* and the Δ*pilY1* mutant *(C)* on glass and bulk agarose gel surfaces, determined by plate counting method. The first timepoint shown, time = 0 h, occurs after bacteria have been allowed to accumulate to surfaces for one hour. Replicate experiments are indicated by −1 and −2 on the same surface type. Color blocks show the average length of the lag phase on glass (blue) and bulk gel (orange) surfaces.

**Fig. S3.** The configuration change *(A)* and cell volume change *(B)* as a cell envelope attaches to stiff and soft substrates using finite element models. The mesh lines are rendered and the configurations at a displacement of 150 nm are shown. Cell volume is normalized to that in the free-floating state. *(C)* Approximating bacterial surface adhesion by displacing a cell envelope toward substrates. The schematic illustration of the adhesion force scheme for modeling surface adhesion. Arrows denote the direction of adhesion forces, and the highlighted area denotes the area over which the forces are applied. Note that the same forces with the opposite direction are applied on the substrate contact surface but they are not shown here for brevity. *(D)* Comparing the displacement and adhesion force schemes. The circumferential stresses at element #1 with different loading schemes are compared. The dash line denotes when the cell first contacts the substrate. CW: cell wall; OM: outer membrane.

**Fig. S4.** The spatial distribution of outer membrane stresses *(A-C)* and contact pressure *(D-F)*. Stresses are normalized to their respective values during the free-floating state. The dash line denotes when the cell first contacts the substrate. *(G and H)* Mechanical strain state on the inner membrane, and its spatial distribution *(I)*. The strain distribution is illustrated with a model where the substrate is stiff and the displacement is 150 nm. Note that the strain here is calculated based on a stress-free reference state. The mesh lines are hidden and the strains in the substrate are not shown. (*J*) Convergence studies of the finite element models. The circumferential stress of the outer membrane at element #1 is compared at different mesh sizes. In all images, subscript c denotes the circumferential direction and subscript a denotes the axial direction.

**Fig. S5.** Bacterial surface motility during the first one hour of the accumulation process. *(A)* Among all tracked bacteria on thin or thick agarose gel composites, the percentage of WT and the Δ*pilY1* mutant remaining stationary and showing flagellum-driven spinning motility or TFP-driven twitching motility. *(B)* The percentage of spinning or twitching WT and the Δ*pilY1* mutant, accounting for all motile bacteria attached to surfaces. *(C)* Detachment of adhering WT and the Δ*pilY1* mutant from thin and thick agarose gel surfaces during the first one hour of the accumulation process. WT are significantly more likely to detach from thick gels (30 detachment events among 673 tracked cells) than from thin gels (10 detachment events among 1609 tracked cells) (*P* <0.001, χ^2^ test), while Δ*pilY1* are equally likely to detach from thin and thick gels (*P* = 0.78, χ^2^ test). *(D)* The linear correlation for the median value of bacteria speed. *(E)* The linear correlation for the mean value of bacteria speed. *** *P* <0.001; Mann-Whitney u test. * *P* <0.05; based on the 95% confidence interval of mean value of bacteria speed reported in the Main text. These indicate that bacteria speed of the Δ*pilY1* mutant on gels is significantly higher than that of WT on thin gels, but significantly lower than that of WT on thick gels, for both median value and mean value of bacteria speed.

**Fig. S6.** *(A* and *B)* The corresponding relations between the mean fluorescence intensity of beads attached on thick agarose gels and on thin agarose gels. *(A)* The relation for intracellular Na^+^ measurement, in which beads were imaged within NaCl buffer in an imaging spacer (0.12 mm depth). *(B)* The relation for c-di-GMP measurement, in which beads were imaged within LB medium in an imaging chamber (2.6 mm depth). The exponential equation is the fitted curve to the dataset. Growth dynamics of attached Δ*pilY1*/P_*BAD*_∷*pilY1 (C)* and Δ*pilT*/P_*BAD*_∷*pilT* complements *(D)* on thin and thick agarose gel composites. Data are means ± SD. The data at 0 time point corresponds to the end of one hour of bacterial accumulation on gel surfaces.

**Fig. S7.** Phase contrast images of WT *(A and D)* and the *ΔpilT (B and E)* and *ΔpilY1 (C and F)* mutants after six hours of incubation on thin and thick agarose gels. The inset in *(B)* shows a magnified image of the upper layer of cells in a micro-colony cluster of Δ*pilT*, in the area indicated by the dotted box.

**Table S1.** Physical/mechanical properties in the finite element models.

**Table S2**. Strains, plasmids and primers used in this study.

## References

1. Cheng B, Lin M, Huang G, Li Y, Ji B, Genin GM, Deshpande VS, Lu TJ, Xu F. 2017. Cellular mechanosensing of the biophysical microenvironment: a review of mathematical models of biophysical regulation of cell responses. Phys Life Rev 22:88–119.

2. Iskratsch T, Wolfenson H, Sheetz MP. 2014. Appreciating force and shape—the rise of mechanotransduction in cell biology. Nat Rev Mol Cell Biol 15:825–833.

3. Persat A. 2017. Bacterial mechanotransduction. Curr Opin Microbiol 36:1–6.

4. Gordon VD, Wang L. 2019. Bacterial mechanosensing: the force will be with you, always. J Cell Sci 132:jcs227694.

5. Dufrêne YF, Persat A. 2020. Mechanomicrobiology: how bacteria sense and respond to forces. Nat Rev Microbiol :1–14.

6. Ellison CK, Kan J, Dillard RS, Kysela DT, Ducret A, Berne C, Hampton CM, Ke Z, Wright ER, Biais N. 2017. Obstruction of pilus retraction stimulates bacterial surface sensing. Science 358:535–538.

7. Hug I, Deshpande S, Sprecher KS, Pfohl T, Jenal U. 2017. Second messenger–mediated tactile response by a bacterial rotary motor. Science 358:531–534.

8. Persat A, Inclan YF, Engel JN, Stone HA, Gitai Z. 2015. Type IV pili mechanochemically regulate virulence factors in *Pseudomonas aeruginosa*. Proc Natl Acad Sci USA 112:7563–7568.

9. Luo Y, Zhao K, Baker AE, Kuchma SL, Coggan KA, Wolfgang MC, Wong GC, O’Toole GA. 2015. A hierarchical cascade of second messengers regulates *Pseudomonas aeruginosa* surface behaviors. mBio 6:e02456–14.

10. Siryaporn A, Kuchma SL, O’Toole GA, Gitai Z. 2014. Surface attachment induces *Pseudomonas aeruginosa* virulence. Proc Natl Acad Sci USA 111:16860–16865.

11. Talà L, Fineberg A, Kukura P, Persat A. 2019. *Pseudomonas aeruginosa* orchestrates twitching motility by sequential control of type IV pili movements. Nat Microbiol 4:774–780.

12. O’Neal L, Baraquet C, Suo Z, Dreifus JE, Peng Y, Raivio TL, Wozniak DJ, Harwood CS, Parsek MR. 2022. The Wsp system of Pseudomonas aeruginosa links surface sensing and cell envelope stress. Proc Natl Acad Sci USA 119:e2117633119.

13. Rodesney CA, Roman B, Dhamani N, Cooley BJ, Katira P, Touhami A, Gordon VD. 2017. Mechanosensing of shear by *Pseudomonas aeruginosa* leads to increased levels of the cyclic-di-GMP signal initiating biofilm development. Proc Natl Acad Sci USA 114:5906–5911.

14. Sanfilippo JE, Lorestani A, Koch MD, Bratton BP, Siryaporn A, Stone HA, Gitai Z. 2019. Microfluidic-based transcriptomics reveal force-independent bacterial rheosensing. Nat Microbiol 4:1274–1281.

15. Nguyen Y, Sugiman-Marangos S, Harvey H, Bell SD, Charlton CL, Junop MS, Burrows LL. 2015. *Pseudomonas aeruginosa* minor pilins prime type IVa pilus assembly and promote surface display of the PilY1 adhesin. J Biol Chem 290:601–611.

16. Guimarães CF, Gasperini L, Marques AP, Reis RL. 2020. The stiffness of living tissues and its implications for tissue engineering. Nat Rev Mater :1–20.

17. Wang Y, Guan A, Isayeva I, Vorvolakos K, Das S, Li Z, Phillips KS. 2016. Interactions of Staphylococcus aureus with ultrasoft hydrogel biomaterials. Biomaterials 95:74–85.

18. Campoccia D, Montanaro L, Arciola CR. 2006. The significance of infection related to orthopedic devices and issues of antibiotic resistance. Biomaterials 27:2331–2339.

19. Funt D, Pavicic T. 2013. Dermal fillers in aesthetics: an overview of adverse events and treatment approaches. Clin Cosmet Investig Dermatol 6:295.

20. Wald HL, Kramer AM. 2007. Nonpayment for harms resulting from medical care: catheter-associated urinary tract infections. Jama 298:2782–2784.

21. Kolewe KW, Peyton SR, Schiffman JD. 2015. Fewer bacteria adhere to softer hydrogels. ACS Appl Mater Interfaces 7:19562–19569.

22. Peng Q, Zhou X, Wang Z, Xie Q, Ma C, Zhang G, Gong X. 2019. Three-dimensional bacterial motions near a surface investigated by digital holographic microscopy: effect of surface stiffness. Langmuir 35:12257–12263.

23. Kolewe KW, Zhu J, Mako NR, Nonnenmann SS, Schiffman JD. 2018. Bacterial adhesion is affected by the thickness and stiffness of poly (ethylene glycol) hydrogels. ACS Appl Mater Interfaces 10:2275–2281.

24. Song F, Brasch ME, Wang H, Henderson JH, Sauer K, Ren D. 2017. How bacteria respond to material stiffness during attachment: a role of *Escherichia coli* flagellar motility. ACS Appl Mater Interfaces 9:22176–22184.

25. Straub H, Bigger CM, Valentin J, Abt D, Qin XH, Eberl L, Maniura-Weber K, Ren Q. 2019. Bacterial Adhesion on Soft Materials: Passive Physicochemical Interactions or Active Bacterial Mechanosensing? Adv Healthc Mater 8:1801323.

26. Otto K, Silhavy TJ. 2002. Surface sensing and adhesion of *Escherichia coli* controlled by the Cpx-signaling pathway. Proc Natl Acad Sci USA 99:2287–2292.

27. Shimizu T, Ichimura K, Noda M. 2016. The surface sensor NlpE of enterohemorrhagic *Escherichia coli* contributes to regulation of the type III secretion system and flagella by the Cpx response to adhesion. Infect Immun 84:537–549.

28. O’Neal L, Baraquet C, Suo Z, Dreifus JE, Peng Y, Raivio TL, Wozniak DJ, Harwood CS, Parsek MR. 2022. The Wsp system of *Pseudomonas aeruginosa* links surface sensing and cell envelope stress. Proceedings of the National Academy of Sciences 119:e2117633119.

29. Laventie B-J, Jenal U. 2020. Surface sensing and adaptation in bacteria. Annu Rev Microbiol 74:735–760.

30. Booth IR. 2014. Bacterial mechanosensitive channels: progress towards an understanding of their roles in cell physiology. Curr Opin Microbiol 18:16–22.

31. Booth IR, Edwards MD, Black S, Schumann U, Miller S. 2007. Mechanosensitive channels in bacteria: signs of closure? Nat Rev Microbiol 5:431.

32. O’Toole G, Kaplan HB, Kolter R. 2000. Biofilm formation as microbial development. Annu Rev Microbiol 54:49–79.

33. Conrad JC, Gibiansky ML, Jin F, Gordon VD, Motto DA, Mathewson MA, Stopka WG, Zelasko DC, Shrout JD, Wong GC. 2011. Flagella and pili-mediated near-surface single-cell motility mechanisms in *P. aeruginosa*. Biophys J 100:1608–1616.

34. Gibiansky ML, Conrad JC, Jin F, Gordon VD, Motto DA, Mathewson MA, Stopka WG, Zelasko DC, Shrout JD, Wong GC. 2010. Bacteria use type IV pili to walk upright and detach from surfaces. Science 330:197–197.

35. Bennett RR, Lee CK, De Anda J, Nealson KH, Yildiz FH, O’Toole GA, Wong GCL, Golestanian R. 2016. Species-dependent hydrodynamics of flagellum-tethered bacteria in early biofilm development. Journal of The Royal Society Interface 13:20150966.

36. Jenal U, Reinders A, Lori C. 2017. Cyclic di-GMP: second messenger extraordinaire. Nat Rev Microbiol 15:271.

37. Rybtke MT, Borlee BR, Murakami K, Irie Y, Hentzer M, Nielsen TE, Givskov M, Parsek MR, Tolker-Nielsen T. 2012. Fluorescence-based reporter for gauging cyclic di-GMP levels in *Pseudomonas aeruginosa*. Appl Environ Microbiol 78:5060–5069.

38. Laventie B-J, Sangermani M, Estermann F, Manfredi P, Planes R, Hug I, Jaeger T, Meunier E, Broz P, Jenal U. 2019. A surface-induced asymmetric program promotes tissue colonization by *Pseudomonas aeruginosa*. Cell Host Microbe 25:140–152. e6.

39. Bertrand RL. 2019. Lag phase is a dynamic, organized, adaptive, and evolvable period that prepares bacteria for cell division. J Bacteriol 201:e00697–18.

40. Lee CK, de Anda J, Baker AE, Bennett RR, Luo Y, Lee EY, Keefe JA, Helali JS, Ma J, Zhao K. 2018. Multigenerational memory and adaptive adhesion in early bacterial biofilm communities. Proc Natl Acad Sci USA 115:4471–4476.

41. Lee CK, Vachier J, de Anda J, Zhao K, Baker AE, Bennett RR, Armbruster CR, Lewis KA, Tarnopol RL, Lomba CJ. 2020. Social cooperativity of bacteria during reversible surface attachment in young biofilms: a quantitative comparison of *Pseudomonas aeruginosa* PA14 and PAO1. mBio 11.

42. Park S, Sauer K. 2022. Controlling Biofilm Development Through Cyclic di-GMP Signaling, p 69–94. *In* Filloux A, Ramos J-L (ed), Pseudomonas aeruginosa: Biology, Pathogenesis and Control Strategies. Springer International Publishing, Cham.

43. Lichtenberg M, Kragh KN, Fritz B, Kirkegaard JB, Tolker-Nielsen T, Bjarnsholt T. 2022. Cyclic-di-GMP signaling controls metabolic activity in *Pseudomonas aeruginosa*. Cell Reports 41.

44. Webster SS, Mathelié-Guinlet M, Verissimo AF, Schultz D, Viljoen A, Lee CK, Schmidt WC, Wong GC, Dufrêne YF, O’Toole GA. 2022. Force-induced changes of PilY1 drive surface sensing by Pseudomonas aeruginosa. mbio 13:e03754–21.

45. Trivedi RR, Crooks JA, Auer GK, Pendry J, Foik IP, Siryaporn A, Abbott NL, Gitai Z, Weibel DB. 2018. Mechanical genomic studies reveal the role of D-alanine metabolism in Pseudomonas aeruginosa cell stiffness. mbio 9.

46. Formosa C, Grare M, Duval RE, Dague E. 2012. Nanoscale effects of antibiotics on *P. aeruginosa*. Nanomedicine: NBM 8:12–16.

47. Mathelié-Guinlet M, Grauby-Heywang C, Martin A, Février H, Morote F, Vilquin A, Beven L, Delville M-H, Cohen-Bouhacina T. 2018. Detrimental impact of silica nanoparticles on the nanomechanical properties of *Escherichia coli*, studied by AFM. J Colloid Interface Sci 529:53–64.

48. Rojas ER, Billings G, Odermatt PD, Auer GK, Zhu L, Miguel A, Chang F, Weibel DB, Theriot JA, Huang KC. 2018. The outer membrane is an essential load-bearing element in Gram-negative bacteria. Nature 559:617–621.

49. Tuson HH, Auer GK, Renner LD, Hasebe M, Tropini C, Salick M, Crone WC, Gopinathan A, Huang KC, Weibel DB. 2012. Measuring the stiffness of bacterial cells from growth rates in hydrogels of tunable elasticity. Mol Microbiol 84:874–891.

50. Thwaites JJ, Mendelson NH. 1991. Mechanical Behaviour of Bacterial Cell Walls, p 173–222. *In* Rose AH, Tempest DW (ed), Advances in Microbial Physiology, vol 32. Academic Press.

51. Timoshenko SP, Woinowsky-Krieger S. 1959. Theory of plates and shells. McGraw-hill.

52. Matias VR, Al-Amoudi A, Dubochet J, Beveridge TJ. 2003. Cryo-transmission electron microscopy of frozen-hydrated sections of Escherichia coli and *Pseudomonas aeruginosa*. J Bacteriol 185:6112–6118.

53. Hayhurst EJ, Kailas L, Hobbs JK, Foster SJ. 2008. Cell wall peptidoglycan architecture in Bacillus subtilis. Proc Natl Acad Sci USA 105:14603–14608.

54. Auer GK, Weibel DB. 2017. Bacterial Cell Mechanics. Biochemistry 56:3710–3724.

55. Vadillo-Rodriguez V, Schooling SR, Dutcher JR. 2009. In situ characterization of differences in the viscoelastic response of individual gram-negative and gram-positive bacterial cells. J Bacteriol 191:5518–5525.

56. Kumar U, Vivekanand K, Poddar P. 2009. Real-time nanomechanical and topographical mapping on live bacterial cells-*Brevibacterium casei* under stress due to their exposure to Co^2+^ ions during microbial synthesis of Co_3_O_4_ nanoparticles. J Phys Chem B 113:7927–7933.

57. Francius G, Domenech O, Mingeot-Leclercq MP, Dufrêne YF. 2008. Direct observation of *Staphylococcus aureus* cell wall digestion by lysostaphin. J Bacteriol 190:7904–7909.

58. Saha N, Monge C, Dulong V, Picart C, Glinel K. 2013. Influence of polyelectrolyte film stiffness on bacterial growth. Biomacromolecules 14:520–528.

59. Guégan C, Garderes J, Le Pennec G, Gaillard F, Fay F, Linossier I, Herry J-M, Fontaine M-NB, Réhel KV. 2014. Alteration of bacterial adhesion induced by the substrate stiffness. Colloids Surf B Biointerfaces 114:193–200.

60. Cloutier M, Mantovani D, Rosei F. 2015. Antibacterial coatings: challenges, perspectives, and opportunities. Trends Biotechnol 33:637–652.

61. Salwiczek M, Qu Y, Gardiner J, Strugnell RA, Lithgow T, McLean KM, Thissen H. 2014. Emerging rules for effective antimicrobial coatings. Trends Biotechnol 32:82–90.

62. Moriarty TF, Zaat SA, Busscher HJ. 2012. Biomaterials associated infection: immunological aspects and antimicrobial strategies. Springer Science & Business Media.

63. Imani SM, Maclachlan R, Rachwalski K, Chan Y, Lee B, McInnes M, Grandfield K, Brown ED, Didar TF, Soleymani L. 2019. Flexible hierarchical wraps repel drug-resistant gram-negative and positive bacteria. ACS Nano 14:454–465.

64. Feng G, Cheng Y, Wang S-Y, Borca-Tasciuc DA, Worobo RW, Moraru CI. 2015. Bacterial attachment and biofilm formation on surfaces are reduced by small-diameter nanoscale pores: how small is small enough? NPJ Biofilms Microbiomes 1:1–9.

65. Matsui H, Wagner VE, Hill DB, Schwab UE, Rogers TD, Button B, Taylor RM, Superfine R, Rubinstein M, Iglewski BH. 2006. A physical linkage between cystic fibrosis airway surface dehydration and *Pseudomonas aeruginosa* biofilms. Proc Natl Acad Sci USA 103:18131–18136.

66. Hutchinson J, Lawrence J. 1991. Wound infection under occlusive dressings. J Hosp Infect 17:83–94.

67. Jacobs MA, Alwood A, Thaipisuttikul I, Spencer D, Haugen E, Ernst S, Will O, Kaul R, Raymond C, Levy R. 2003. Comprehensive transposon mutant library of *Pseudomonas aeruginosa*. Proc Natl Acad Sci USA 100:14339–14344.

